# Optimizing Genomic Data Compression with Genetic Algorithms

**DOI:** 10.1101/2025.10.23.684090

**Authors:** Rita Ferrolho, Armando J. Pinho, Diogo Pratas

## Abstract

The rapid growth of genomic data, especially in clinical settings, has highlighted the need for efficient lossless compression. Preserving full sequence information is critical, as even small data losses can affect analyses or diagnoses. With increasing dataset size and complexity, optimized compression is vital for storage, transmission, and processing. Fixed-parameter compression tools often fall short across diverse genomes, demonstrating the need for adaptive tuning strategies tailored to each dataset. In this paper, we investigate the use of metaheuristic search algorithms, particularly Genetic Algorithms (GAs), to optimize parameter configurations for JARVIS3, a genomic sequence compression tool. Due to JARVIS3’s large and complex parameter space, exhaustive search is infeasible. To tackle this, we developed OptimJV3, a modular framework that integrates multiple GA variants, including a multi-objective approach that also considers computational time. We evaluated OptimJV3 on various genomic datasets (FASTA format), namely Human Chromosome Y (CY), Cassava, and the full Human Genome (HG), and observed notable compression gains. For large sequences like Cassava and HG, we applied sampling-based optimization: optimal parameters were first identified from smaller segments and then used for full sequence compression. These small segments proved effective for guiding parameter tuning. Further improvements were achieved by increasing the number of GA generations, especially for HG. For example, using a 10 MB sample and 500 generations, OptimJV3 reached a compression rate of 1.431 bits per base on HG - among the best results reported so far.

## 1 Introduction

The exponential increase in genomic data production, driven by advances in high-throughput sequencing (HTS) technologies, has introduced significant challenges in data storage, transmission, and computational analysis. This issue is particularly critical in the clinical and biomedical research domains, where lossless compression is essential to ensure the integrity of sequence data. Even minor information loss can lead to incorrect diagnostic interpretations or flawed downstream analyses [56].

Although state of the art tools like JARVIS3 have been developed to compress genomic sequences efficiently, a major limitation is their reliance on fixed parameter configurations, which may not produce optimal results across diverse genomic datasets [60]. This is especially problematic given the variability in genomic structure, repeat content, and organism origin. Additionally, the parameter space of JARVIS3 is vast and complex, making exhaustive or manual searches for optimal settings infeasible.

An important feature is that, in many clinical applications, compression and decompression workflows are performed repeatedly on versions of the genome that differ only slightly from the reference. In fact, more than 95% of genomic data in such cases is identical, with only a small proportion of variation (e.g. SNPs, indels). As a result, tuning the compression parameters for a specific reference genome is typically a one-time process that can deliver long-term benefits through the repeated use of optimized settings.

Therefore, we hypothesize that metaheuristic algorithms, particularly Genetic Algorithms (GAs), offer a promising solution for this optimization challenge. These methods are designed to efficiently navigate large, nonlinear, and multidimensional search spaces and are well-suited for identifying high-quality parameter configurations that maximize compression performance while minimizing computational cost. Moreover, multi-objective optimization enables simultaneous consideration of compression ratio and processing time, an essential trade-off in real-world scenarios.

To test this hypothesis, we developed a new method along with its implementation, OptimJV3, which is a modular and extensible optimization framework based on GAs. The tool supports multiple GA variants and integrates multi-objective search strategies. We applied OptimJV3 to optimize JARVIS3 compression settings in several genomic datasets, including Human Chromosome Y (CY), Cassava, and the entire Human Genome (HG). For large genomes, we introduced a sampling-based optimization strategy, where small sequence segments are used to identify optimal parameters that are later applied to the full dataset. This approach significantly reduces the optimization time while achieving strong compression performance in different genome sizes and types.

A crucial foundation for understanding the need for optimization lies in prior research on biological data compression methods [17], which traces back to the development of Biocompress [14]. This early algorithm identifies repetitive sequences and complementary structures, such as palindromes, encoding them based on their length and the position of their earliest occurrence. In specific genomic contexts, Biocompress demonstrated superior performance compared to conventional compression approaches.

Since Biocompress several DNA compression algorithms have been developed over the years to handle the unique structure and patterns of genetic sequences. Some of these algorithms include *Cfact* [15], *CDNA* [47], *ARM* [1], *Offline* [2], *DNACompress* [8], *CTW + LZ* [28], *NMLComp* [69], *GeNML* [18], *DNASequitur* [9], *DNA-X* [26], *DNAC* [20], *XM* [3], XM [5], *Differential Direct Coding (2D)* [71], *DNASC* [29], *GBC* [46], *DNACompact* [16], *POMA* [75], *DNAEnc3* [31], *GenCodex* [52], *BIND* [4], and *DNA-COMPACT* [21], HighFCM [37], SeqCompress [51], CoGi [73], GeCo [42], OCW [7], OBComp [25], DeepDNA [72], JARVIS [36], GeCo2 [35], GeCo3 [59], JARVIS2 [41], Gencoder [54], DNACoder [53], GC-2 [55], LEC-Codec [68], GMG [30], TARG [48], GraSS [49], AGDLC [62], JARVIS3 [60]. Additionally, to enable the compression of both headers and DNA sequences (FASTA format), several specialized tools have been developed, including MFCompress [32], MemRGC [22], NAF [19], MBGC [13], and AGC [12].

Beyond domain-specific genome compressors, recent work explores lossless compression based on learning, often universal or multi-source [63]; TRACE uses transformers for fast general-purpose coding [27]; and MSDZip plus an xLSTM-mapping variant target universal/multi-source settings [23, 24]. For genomics, LRCB benchmarks reference-free long-read tools [65], while PMKLC, (S,K)-mer + DNN, PMFFRC, and SR2C introduce learning/structure-aware methods for large repositories or short-read redundancy [61, 66, 67, 64].

From several benchmarks (e.g. [10] and [11]), we can see that JARVIS3 [60] provides, on average, the highest compression results while using balanced computational resources. Specifically, JARVIS3 is an efficient reference-free genomic data compressor designed to meet the storage demands of large-scale genomic projects. JARVIS3 uses a cooperation between finite-context models (CMs) [43] and weighted stochastic repeat models (WSRMs or RMs) [33].

Specifically, CMs are statistical models that rely on the Markov property. An order-k CM assigns probability estimates to symbols in the alphabet based on a conditioning context derived from a fixed number (k) of preceding outcomes [31].

In contrast, an RM predicts the next symbol by identifying a match with a previously observed symbol. It maintains a pointer to the position of the symbol being copied, along with auxiliary information used to assess the reliability of its predictions [6, 34].

The cooperation within and between CMs and RMs can be provided using a soft-blending approach or a neural network. It offers three profiles, from rapid to high-compression modes, to suit various computational needs, and its flexible C/Bash implementation supports FASTA and FASTQ data as well as parallel processing.

Our benchmarks and its flexibility indicate that JARVIS3 is among the strongest reference-free compressors for assembled genomes, delivering competitive (often superior) BPS (bits per symbol) at balanced runtime. As described, its architecture combines many distinct models and powerful compositions (See Supplementary Subsection 5.7 JARVIS3 Parameters), enabling state-of-the-art results; however, this flexibility creates a large, high-dimensional parameter space. OptimJV3 addresses this need by automatically searching and tuning these parameters, resulting in consistent gains without manual effort.

In this paper, we explore the use of Genetic Algorithms (GAs) for parameter optimization in JARVIS3 [60]. Our goal is to identify configurations that enhance both compression ratio and processing speed specifically for reference genomes or genome collections (e.g., pangenomes) in FASTA format. For this purpose, we introduce OptimJV3, an open-source optimization framework that employs multiple variants of Metameric Genetic Algorithms (MGAs) to systematically tune JARVIS3 for various genomic sequences.

The remainder of this paper is organized as follows: we first describe the optimization algorithm in detail, then present and discuss our experimental results and, finally, conclude with insights and future directions.

## 2 Method

The proposed method (OptimJV3) in this paper is based on a MGA framework that searches for optimal JARVIS3 models and parameters to compress a given sequence. A GA is considered metameric when the algorithm operates on individuals with a metameric structure, that is, a body divided into a series of structurally similar, but not necessarily identical, segments [50].

The MGA begins by generating an initial population of individuals, that is, JARVIS3 commands, and then iterates through several key operations to evolve the population: run, evaluation, selection, crossover, and mutation. Figure 1 illustrates the overall method used by OptimJV3.

**Figure 1.**
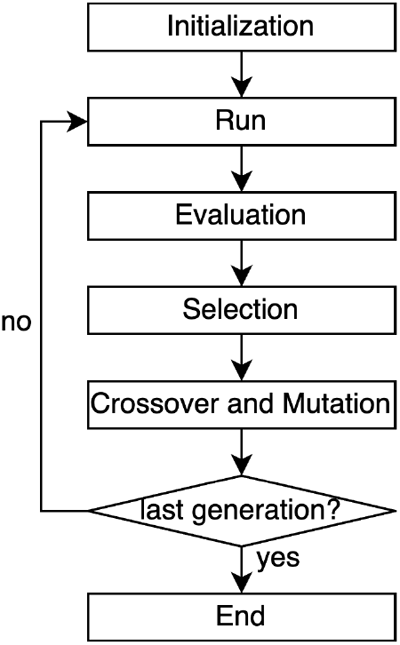
Overview of the OptimJV3 method while using a Genetic Algorithm.

The algorithm ends when the last generation, defined by the user or the program, is reached. Furthermore, individuals within a population may possess varying numbers of CMs and RMs, and the implemented MGA is capable of functioning effectively on individuals exhibiting such diversity. An example of this structure is shown in A) from Figure 2.

**Figure 2.**
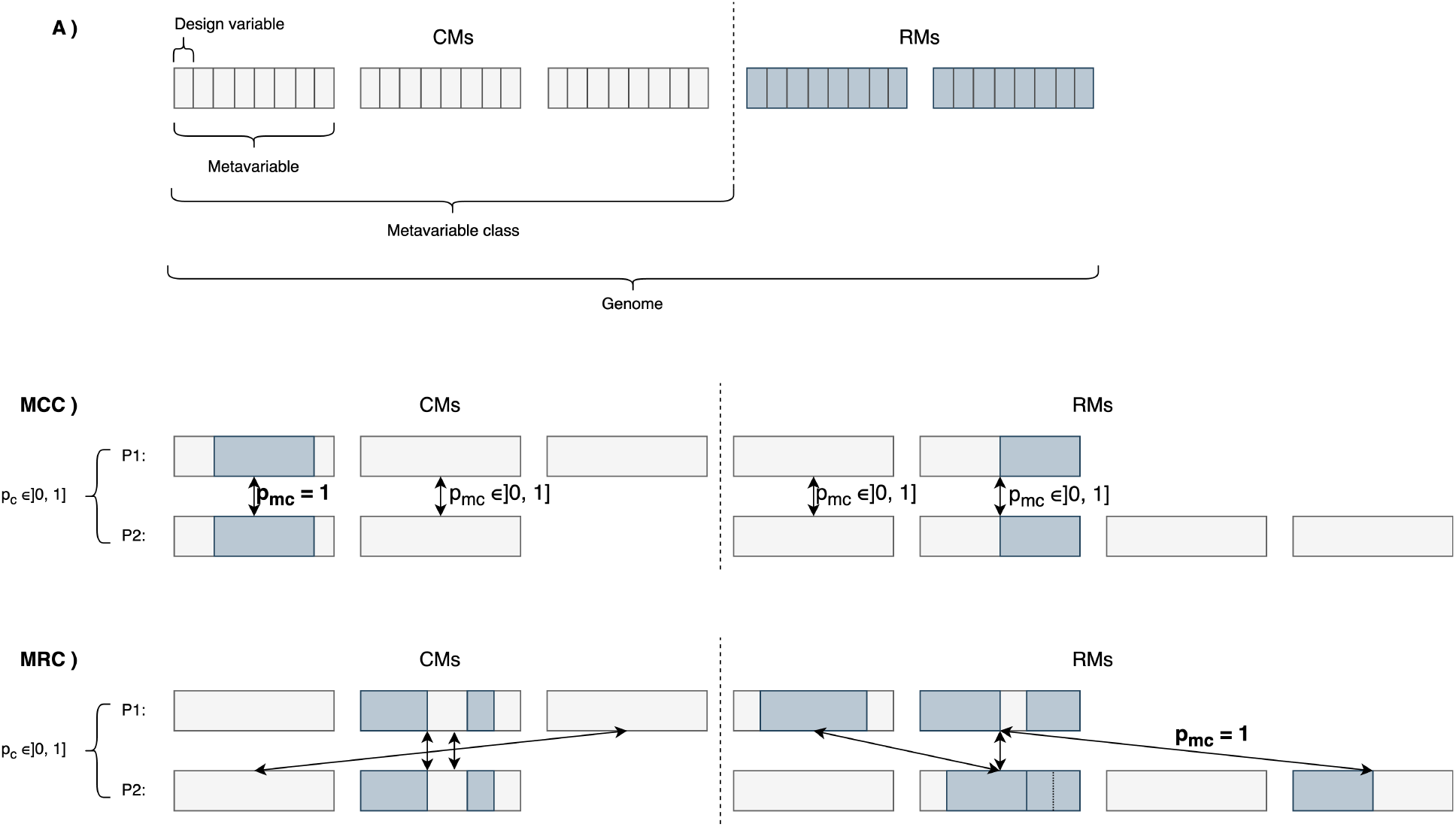
A) Metameric structure of a JARVIS3 instruction containing three context models (CMs) and two repeat models (RMs); MCC) Metameric Canonical Crossover, where metavariable pairs are formed by systematically matching corresponding metavariables from each parent, until all metavariables from the shorter parent are paired. A crossover operator is then applied to each pair selectively, based on a given probability; MRC) Metamerical Random Crossover operates similarly to MCC, but the metavariable pairs are chosen randomly. A metavariable may appear in more than one pair, but pairs composed of metavariables from the same parent are not allowed.

The optimization process begins with an initial population of candidate solutions, which can be generated randomly, where CM and RM parameters of each individual are generated within a broader search area; heuristically, where these parameters are generated within a narrower search area; or in hybrid mode, where some individuals are generated randomly and others heuristically. All initialization methods generate a population that can evolve with a dynamic number of CMs and RMs in each individual. By default, each individual contains 1 to 5 CMs and 1 to 4 RMs, independently of the generation. Although these constraints are enforced by default, adjusting these values within the program is straightforward.

Several search strategies are used in this study, namely random search (RS), local search (LS), and MGA. RS samples feasible commands uniformly at random and evaluates them independently, while providing breadth but without learning from past trials. LS starts from a seeded command and performs greedy or hill-climbing moves in a small neighborhood of edits; it can quickly refine good seeds but is sensitive to initialization and can get stuck in local optima. Our MGA is population-based, meaning that it evolves variable-length (“metameric”) command strings via selection, crossover, and mutation, maintaining diversity while optimizing single- or multi-objective fitness (e.g., compression vs. time). MGA tends to explore globally and discover efficient commands at higher compute cost per generation.

After running the individuals, they are evaluated based on their fitness score, which is determined by multiple criteria, including validity, the size of the compressed sequence, and compression time. The fitness function can operate under either a single-objective (SOGA) or a multi-objective (MOGA) framework, with the latter allowing for weighted metrics to balance the importance of different objectives during optimization, namely compression ratio and computational time. By default, a SOGA fitness function, ℱ, is applied according to

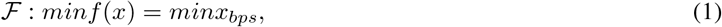

where *x*_*bps*_ represents the BPS compression. If the MOGA feature is chosen, the fitness scores can be calculated through a weighted metric function defined as

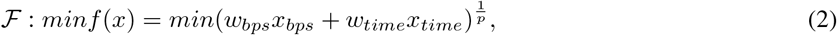

where *x*_*time*_ corresponds to the compression time (seconds) and *w*_*time*_ represents its respective weight, *x*_*bps*_ corresponds to the compression value in BPS and *w*_*bps*_ represents its respective weight, and *p* is a variable with default value of 2. The default weight values are *w*_*time*_ = *w*_*bps*_ = 0.5.

The next step is selection, which involves identifying the fittest individuals from the population, with options for various selection methods such as elitism, modified roulette wheel selection (RWS), and tournament selection [74]. The modified RWS is identical to the standard RWS, but each individual can only be selected once. This is achieved by removing the selected individual from the roulette and recalculating the probabilities of each remaining individual before selecting the next.

After selection, individuals are paired for crossover, which combines their genetic information to produce new offspring based on specified crossover rates applied to a pair of genomes (default rate: 0.6) or metavariables (default rate: 0.6). Two metameric crossover operators can be used: the metameric canonical crossover (MCC) and metameric random crossover (MRC), tailored to the specific structures of command and model parameters, as shown in Fig. 2. The available crossover operators applied to a pair of CM or RM models (default rate: 0.6) are x-point, uniform, average, discrete, flat, and heuristic [70].

Finally, mutation may occur, where selected offspring undergo random changes in one or more of their parameters, ensuring diversity and preventing premature convergence of the population. This mutation can be either heuristic, where the value of a chosen parameter of a CM or RM is changed within a narrower range of possible values; or random, where this value is changed within a broader range. By default, each individual has a mutation rate of 0.1, and assuming the individual is mutated, each parameter has a default mutation rate of 0.1.

Notice that using a GA to search OptimJV3’s parameters is practical because it handles mixed variable types, hard constraints, and variable-length command structures, and it is robust to non-convex or noisy objectives while supporting multi-objective trade-offs (compression vs. time) and easy parallelism. Limitations include higher evaluation cost than model-based methods, sensitivity to meta-parameters, and potential premature convergence without diversity controls. By contrast, Bayesian optimization is often more sample-efficient on low-dimensional, smooth, well-constrained spaces, and neural/surrogate models are fast after training but need substantial labeled data and retraining to generalize. Overall, GA offers a simple, domain-agnostic search engine for OptimJV3 with clear trade-offs.

### 2.1 Implementation

We implemented the proposed method as a computational tool named OptimJV3. The tool supports user -defined parameters - including population size, genetic operators, and their corresponding rates—offering flexibility and adaptability across diverse optimization scenarios. Additionally, OptimJV3 generates evolution plots and histograms, both for individual sequences and for groups of sequences, facilitating comprehensive performance analysis.

OptimJV3 is implemented in Bash and integrates several external tools, which can be installed by executing the InstallTools.sh script. The complete source code, along with scripts for reproducing the results, is freely available at the project repository: https://github.com/cobilab/OptimJV3. Comprehensive documentation, including detailed instructions and menu descriptions, is provided in the Supplementary Material. Additionally, the repository’s README file includes brief usage examples to guide users. The software is distributed under the GNU General Public License (GPLv3), and full licensing information is available within the repository.

## 3 Results and Discussion

The use of OptimJV3 for optimizing JARVIS3 commands yielded significant improvements in both compression size and execution time for the data sequences tested. The algorithm demonstrated its effectiveness in navigating a diverse search space by employing a steady-state approach that facilitated the continuous evolution of populations. Specifically, we applied several optimization methods, namely RS, LS, MGAs, and sampling, to enhance genomic sequence compression.

For context, RS provides breadth without feedback, LS fine-tunes from seeds within a local neighborhood, and MGA balances global exploration and exploitation via population-based evolution, which helps interpret the trends reported below.

Moreover, unless stated otherwise, reported “Time” refers to the compression runtime of the final command only; the GA search/parameter-tuning overhead is tracked separately and is not included in these numbers.

### 3.1 Datasets and materials

All tested datasets are detailed in the Supplementary Material. Most of them were downloaded from National Center of Biotechnology Information (NCBI) or from a DNA Corpus that includes a set of processed sequences downloaded from NCBI [40]. A few sequences, particularly Cassava haplotigs 1 and 2, were downloaded from the China National GeneBank (CNGB). Additionally, three synthetic sequences of different sizes and patterns were generated using the AlcoR tool [58].

All datasets used in this study are described in Supplementary Table S1, including additional sequences not detailed in the main text that were used for supplementary benchmarking.

All mentioned sequences underwent parameter optimization for compression using RS and LS methods. Additionally, the sequences from Escherichia coli (E. coli), Human chromosome Y (CY), Cassava Haplotig 1 (Cassava), and HG were also optimized using MGA-based methods.

### 3.2 Non-MGAs experiments

Non-MGA approaches, including RS and LS, demonstrated varying success based on dataset size. For sequences under 1 MB, RS achieved fast compression times, with some results matching benchmark compressors in both speed and effectiveness. However, on larger datasets, RS encountered limitations for sequences with dimensions similar or greater than those of Cassava haplotig 2 (approximately 674 MB), and LS found limiting results for sequences with equal or greater length than Panthera Leo (approximately 2 GB), as evidenced in Figure S1 from the Supplementary Material. This highlighted the need for more sophisticated algorithms to tackle extensive datasets and improve reliability and resource demands.

### 3.3 Canonical MGA experiment

In this study, we also addressed the use of a Canonical Metameric Genetic Algorithm (CMGA) specifically for compressing genomes, with tests on Escherichia coli (E. coli), Human chromosome Y (CY), and Cassava haplotig 1 (Cassava). The CMGA is defined as an algorithm that starts a population with 100 individuals, where half of them are generated randomly and the other half heuristically; sorts the results in ascending order by BPS, followed by time efficiency; uses MCC with 0.6 rate and X-point crossover with 0.6 rate; followed by mutation of individuals with rate 0.1.

As depicted in Fig. 3, after several generations, the best-performing individuals achieved substantial reductions in compressed sequence sizes, often exceeding initial benchmarks established during the heuristic initialization phase. The CMGA yielded promising results, especially with CY, which demonstrated rapid convergence and high compressibility. However, the CMGA faced premature convergence, especially with CY. Smaller sequences, such as E. coli and CY, benefited from varied CM parameters, whereas Cassava proved more challenging, pointing to the need for further adjustments to balance computational costs with compression gains. These results are not entirely unexpected, in spite of E. coli sequence length being smaller, as the entropy of this sequence is higher than CY, thus less compressible. Since Cassava has a substantially greater length than E. coli and CY, its optimization required stronger memory resources and time, thus its search was interrupted after a few generations. To be able to optimize sequences with similar or greater length than Cassava, alternative methods were explored.

**Figure 3.**
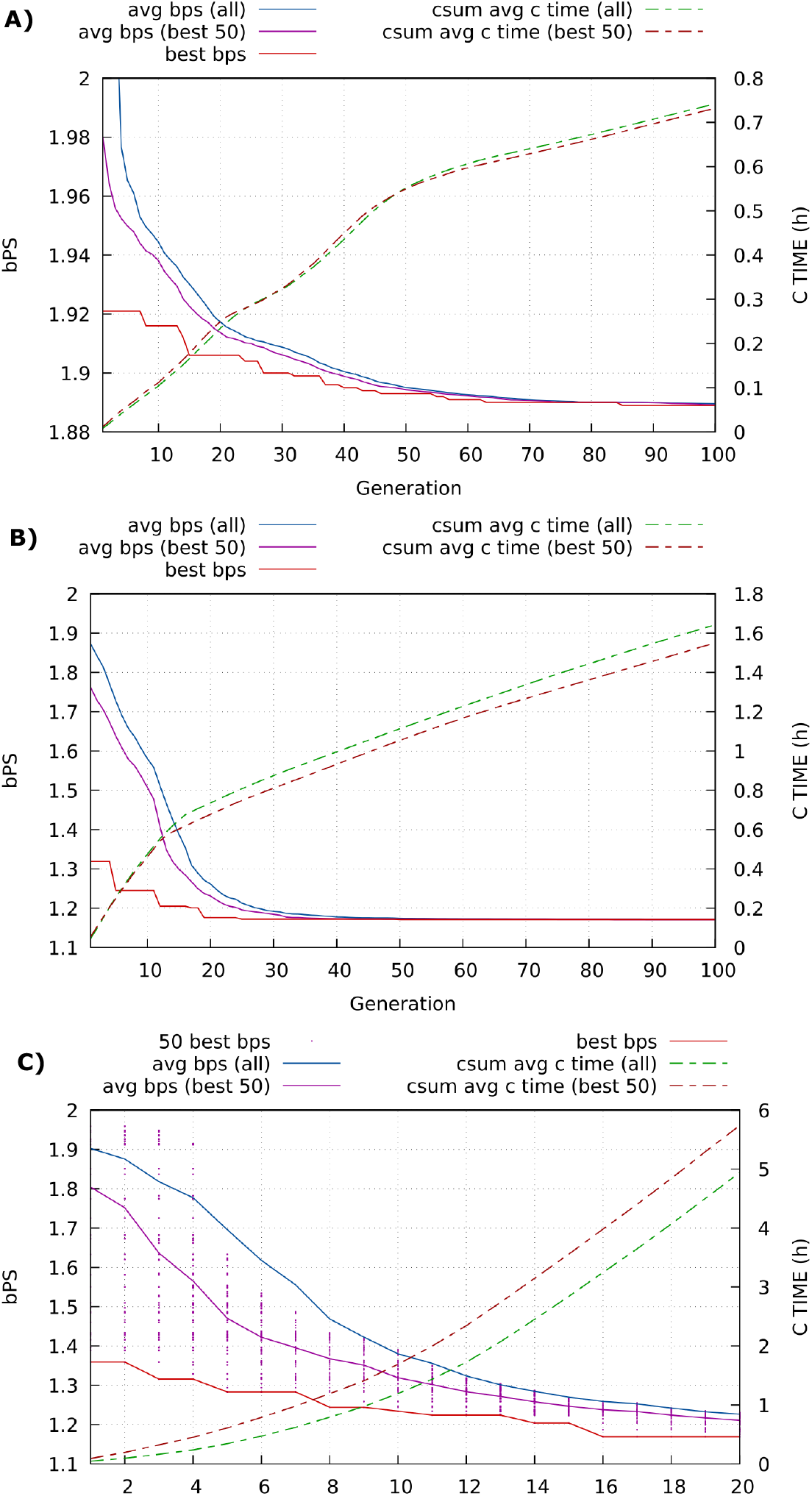
Performance in bits per symbol (BPS) and time (hours) of OptimJV3 for different generations in the optimization of A) Escherichia coli (E. coli, 4.4 MB); B) Human chromosome Y (CY, 21.6 MB); and C) Cassava haplotig 1 (Cassava, 727.0 MB). The csum and avg stands for cumulative sum and average, respectively.

### 3.4 CY experiments

Several MGAs methods were tested and applied to the CY sequence. These methods were organized into five distinct experiments, each exploring different operators and variable adjustments - initialization, population size, evaluation, selection, crossover. The comparison of these methods are illustrated in Fig. 4.

**Figure 4.**
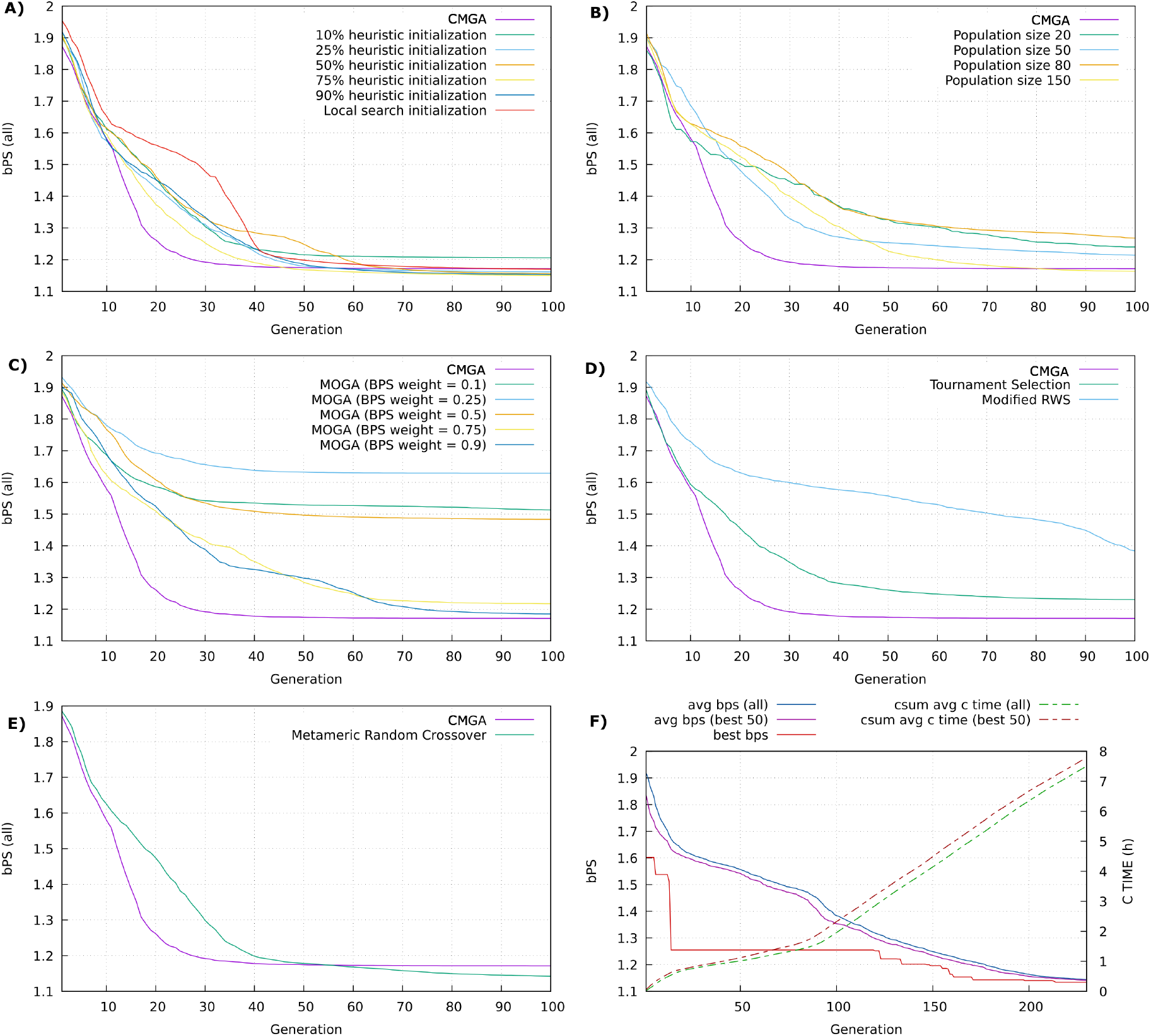
Comparison of several MGA methods applied to CY compression optimization, where y-axis is the BPS average per generation and A) comparison of initialization operators; B) comparison of population sizes; C) comparison of evaluation methods; D) comparison of selection operators; E) comparison of metameric crossover operators. F) is the evolution plot of MGA applied to CY, using modified RWS, from generation 1 to 230.

In the first experiment, different initialization methods, namely random/global, heuristic/local, and hybrid, were tested, revealing that heuristic approaches, though slower to converge, provided more robust solutions than random initialization.

As demonstrated in the second experiment, the population size emerged as a critical factor in the MGA’s performance, particularly for compressing Human chromosome Y. Larger populations increased computational demands. However, a population size of 80 struck a good balance, achieving both competitive compression efficiency and improved processing times. The CMGA, with a population size of 100, ultimately produced the best overall results in terms of compression ratio and speed. This configuration highlighted the MGA’s ability to balance between effective compression and manageable processing time.

The tests on the MOGAs to optimize both the compression ratio and time were successful. These MOGAs converged slower than single-objective MGAs, indicating a strong potential for time optimization, although the results for the compression ratio varied based on weight settings. Among selection and crossover techniques, modified RWS allowed broader search exploration but required more generations to properly converge, while MRC achieved optimized compression values with slower convergence than MCC.

In the fourth experiment, the CMGA, which uses elitist selection, was compared to the tournament and the modified RWS operators. The modified RWS provided the slowest convergence, followed by tournament selection, then elitist selection. These observations are unsurprising, considering that elitist selection generally allows fewer possible selections than other operators, leading to faster convergence. Considering that the modified RWS did not converge after 100 generations, it was decided to proceed the optimization with this operator until it was near convergence, which became more noticeable between generations 200 and 230.

Finally, in the last experiment, the crossover operator used in CMGA, MCC, and the MRC operator were compared. MRC had a slower convergence and reached more optimal BPS results than MCC. These results were expected since the MCC operator presents fewer possible combinations of model pairs than the MRC operator.

Table 1 lists, in ascending order, the best BPS achieved by the evaluated algorithms, and Table 2 reports the fastest solutions. MOGA’s strength is most evident in Table 2—it is the fastest method—highlighting the potential of MOGAs to deliver high-speed solutions or to balance compression efficiency with speed.

**Table 1:** Lowest BPS result returned by some different methods applied to the optimization of CY compression. Time is in seconds and Memory in GigaBytes.

| Rank | Algorithm | BPS | Time | Memory |
| --- | --- | --- | --- | --- |
| 1 | RWS (over 100 generations) | 1.134 | 135.04 | 1.864 |
| 8 | MRC | 1.139 | 129.89 | 2.48 |
| 142 | hybrid initialization | 1.145 | 38.00 | 2.727 |
| 294 | heuristic initialization | 1.149 | 54.26 | 2.505 |
| 2388 | population size 150 | 1.155 | 67.68 | 1.484 |
| 3944 | CMGA | 1.171 | 28.01 | 0.267 |
| 4995 | MOGA (weight BPS = 90%) | 1.180 | 19.97 | 1.677 |

**Table 2:** Fastest result returned by some different methods applied to the optimization of CY compression. Time is in seconds and Memory in GigaBytes.

| Rank | Algorithm | BPS | Time | Memory |
| --- | --- | --- | --- | --- |
| 1 | hybrid initialization | 1.897 | 1.69 | 0.008 |
| 3 | MOGA (weight BPS = 90%) | 1.904 | 1.78 | 0.008 |
| 8 | RWS | 1.897 | 1.85 | 0.008 |
| 10 | heuristic initialization | 1.897 | 1.86 | 0.008 |

Notice that in Tables 1 and 2, “Rank” is the position of an evaluated candidate among all candidates generated during the corresponding GA run (sorted by BPS, then time); e.g., a maximum rank of 4,995 means 4,995 candidates were evaluated.

### 3.5 Cassava sampling experiment

A sampling method has been applied to samples of several sizes of the Cassava sequence - 12.5 MB, 25 MB, 50 MB, and 100 MB - to optimize their compression. To achieve this, a segment of 100 MB, starting from 40% of the sequence (to avoid telomeric regions), was saved as a separate sequence file. The samples of the first 50 MB, 25 MB, and 12.5 MB of the 100 MB sample were then saved in separate sequence files.

Next, the first 100 generations of the MGA were applied to each sample (SG). To decrease the risk of premature convergence, hybrid initialization was used (50% of the initial population heuristically generated), with half of the individuals being generated in a narrower search area. Additionally, the operators RWS, MRC, and uniform crossover were activated, the rate of selected individuals per generation was increased to 0.4 (default: 0.3), and both crossover rates were raised to 0.7 (default: 0.6). To minimize compression efficiency and compression time simultaneously, a multi-objective weighted metric function was applied with a BPS weight of 0.5. Then, the best solution found for each sample were reapplied to the entire sequence. Results of this optimization are presented in Table 3.

**Table 3:** Results of the Cassava sampling experiment, for samples from size 12.5 MB to 100 MB, for the first 100 generations. The “-cm” and “-rm” options specify the parameters for the context model (CM) and repeat model (RM), respectively. Times is in minutes.

| Size | BPS | Time | Models |
| --- | --- | --- | --- |
| 50 | 0.754 | 63.4 | -cm 12:2750:0:0.40/15:3746:0:0.02<br>-rm 197:13:0.73:3:0.75:1:0.12:4<br>-rm 200:13:0.73:3:0.75:1:0.21:4 |
| 25 | 0.755 | 186.0 | -cm 12:1:0:0.10/2:5:1:0.7<br>-cm 1:2403:0:0.15/2:1:1:0.7<br>-rm 296:13:0.95:11:0.34:1:0.61:3<br>-rm 442:1:0.33:5:0.02:2:0.22:5 |
| 100 | 0.756 | 40.1 | -cm 12:2:1:0.20/0:2:0:0.6<br>-cm 8:200:1:0.70/0:2:0:0.6<br>-rm 50:12:0.99:20:0.45:0:0.13:2<br>-rm 50:13:0.99:20:0.90:2:0.16:4 |
| 12.5 | 0.769 | 61.0 | -cm 13:2000:1:0.30/3:2:1:0.7<br>-cm 13:500:1:0.06/3:4140:0:0.80<br>-cm 4:1:1:0.30/5:2:1:0.1<br>-cm 7:4281:1:0.12/3:4173:0:0.08<br>-rm 100:12:0.95:11:0.70:1:0.21:2<br>-rm 1:13:0.95:11:0.05:2:0.61:2<br>-rm 5:12:0.95:5:0.05:1:0.11:2 |

The best solution, in terms of compression efficiency, was obtained from parameters found in the optimization of the 50 MB sample, while the fastest result was yielded from the 100 MB sample optimization. Smaller samples of 25 MB and 12.5 MB returned non-optimal results.

The optimization of these samples was then carried out from generation 101 to generation 200 (SG200), but with the default selection and crossover rates used in OptimJV3 to speed up convergence. Then, for each sample, the models that resulted in the lowest dominance value were reused in the compression of the whole genome. The results of this optimization are presented in Table 4.

**Table 4:** Results of the Cassava sampling experiment, for samples from size 12.5 MB to 100 MB, for the first 200 generations. The “-cm” and “-rm” options specify the parameters for the context model (CM) and repeat model (RM), respectively. Times is in minutes.

| Size | BPS | Time | Models |
| --- | --- | --- | --- |
| 100 | 0.741 | 37.1 | -cm 1:1:1:0.75/3:2:1:0.9<br>-cm 12:1:1:0.70/0:2:0:0.9<br>-rm 20:13:0.90:7:0.75:1:0.16:3<br>-rm 20:13:0.99:7:0.40:1:0.16:2<br>-rm 5:12:0.90:7:0.75:0:0.31:2 |
| 25 | 0.741 | 86.4 | -cm 1:1:0:0.10/2:5:1:0.7<br>-cm 12:1:0:0.10/2:5:1:0.7<br>-rm 296:13:0.95:5:0.34:1:0.61:3<br>-rm 442:13:0.95:11:0.25:1:0.21:5 |
| 50 | 0.749 | 60.6 | -cm 12:4377:0:0.40/15:3746:0:0.02<br>-rm 200:13:0.73:3:0.46:1:0.12:3<br>-rm 200:13:0.73:3:0.82:1:0.12:4 |
| 12.5 | 0.766 | 71.1 | -cm 12:200:1:0.29/2:2:1:0.9<br>-cm 4:1:1:0.20/5:2:0:0.1<br>-cm 6:500:1:0.85/0:3635:0:0.79<br>-rm 100:12:0.85:5:0.45:1:0.11:1<br>-rm 100:12:0.95:11:0.45:1:0.46:2<br>-rm 100:12:0.95:3:0.73:1:0.39:4 |

Surprisingly, the optimization over small segments yielded a substantial improved compression ratio, demonstrating the efficacy of the approach. Overall, extending the MGA from 100 to 200 generations resulted in further improvements in compression performance and time efficiency.

Furthermore, all results shown in Tables 3 and 4 provide compression efficiency results that surpass most results currently presented in the Cassava benchmark, with reasonable or improved time efficiency. This benchmark is available at https://github.com/cobilab/CassavaGenome.

To clarify user effort on Cassava: although OptimJV3 exposes GA controls, in practice we used a simple, fixed recipe—hybrid initialization with RWS/MRC/uniform crossover and a balanced objective—for the first 100 generations, then explicitly reverted to the default selection and crossover rates when extending to 200 generations to speed convergence. The single best model discovered on the 100 MB sample was then reused unchanged to compress the full 727 MB assembly. This shifts dozens of low-level JARVIS3 choices into an automated, one-time search with only high-level knobs (objective and generation budget) and produces a reusable command.

### 3.6 Human genome sampling experiment

Similarly to the Cassava experiment, we applied OptimJV3 to compress HG segments, optimizing JARVIS3 parameters across various small segment sizes, namely 12.5 MB, 25 MB, 50 MB, and 100 MB, of the HG sample for 100 and 200 generations, which were subsequently used to parameterize the compression of the complete HG. However, the MGA configurations used were used differently from those in the Cassava sampling optimization. For the first 100 generations, those used for the Cassava optimization in the last 100 generations were applied, while those used to optimize Cassava in the first 100 generations were reused in the last 100 generations of HG optimization.

The results of this experiment for 100 and 200 generations are shown in Table 5 and Table 6, respectively.

**Table 5:** Results of the HG sampling experiment, for samples from size 12.5 MB to 100 MB, for the first 100 generations. The “-cm” and “-rm” options specify the parameters for the context model (CM) and repeat model (RM), respectively. Times is in minutes.

| Size | BPS | Time | Models |
| --- | --- | --- | --- |
| 50 | 1.427 | 181.1 | -cm 1:1:0:0.90/3:20:1:0.7<br>-cm 11:50:2:0.90/3:1:1:0.84<br>-rm 100:12:0.80:2:0.10:0:0.26:1<br>-rm 100:13:0.80:4:0.10:1:0.77:3<br>-rm 1:12:0.90:4:0.10:2:0.31:1 |
| 12.5 | 1.438 | 191.1 | -cm 13:500:0:0.55/2:20:0:0.9<br>-cm 3:569:2:0.55/2:20:0:0.9<br>-cm 6:500:1:0.55/2:100:0:0.9<br>-rm 200:13:0.90:7:0.60:0:0.96:2<br>-rm 20:13:0.90:9:0.70:2:0.06:3 |
| 100 | 1.441 | 146.4 | -cm 12:200:0:0.35/1:5:1:0.4<br>-cm 2:10:0:0.55/4:20:0:0.41<br>-cm 3:20:0:0.55/0:2294:0:0.13<br>-rm 50:13:0.71:3:0.65:2:0.66:2<br>-rm 5:13:0.71:3:0.59:1:0.66:1 |
| 25 | 1.443 | 236.2 | -cm 2:5:2:0.45/5:5:0:0.7<br>-cm 3:50:0:0.45/4:20:1:0.1<br>-cm 3:5:2:0.65/1:5:0:0.7<br>-rm 200:12:0.55:2:0.50:0:0.01:2<br>-rm 484:12:0.90:8:0.30:1:0.66:5 |

**Table 6:** Results of the HG sampling experiment, for samples from size 12.5 MB to 100 MB, for the first 200 generations. The “-cm” and “-rm” options specify the parameters for the context model (CM) and repeat model (RM), respectively. Times is in minutes.

| Size | BPS | Time | Models |
| --- | --- | --- | --- |
| 25 | 1.418 | 294.8 | -cm 13:5:1:0.30/1:50:0:0.7<br>-cm 2:5:0:0.45/5:5:0:0.7<br>-cm 3:5:2:0.75/1:2:0:0.7<br>-rm 200:12:0.90:8:0.60:1:0.01:5<br>-rm 200:12:0.90:8:0.60:1:0.01:5 |
| 100 | 1.420 | 213.8 | -cm 12:20:0:0.55/1:2:1:0.6<br>-cm 3:20:0:0.55/0:20:0:0.41<br>-cm 9:2000:1:0.55/0:20:0:0.6<br>-rm 50:13:0.95:14:0.65:1:0.11:2<br>-rm 50:13:0.95:8:0.10:1:0.66:1 |
| 12.5 | 1.422 | 246.6 | -cm 13:2:1:0.95/1:2:0:0.9<br>-cm 2:2:1:0.95/1:2:0:0.9<br>-cm 6:2:0:0.95/2:2:0:0.9<br>-rm 200:12:0.90:6:0.70:1:0.06:4<br>-rm 50:12:0.85:6:0.70:1:0.06:4 |
| 50 | 1.423 | 247.4 | -cm 1:1:1:0.15/1:20:1:0.2<br>-cm 8:4998:1:0.35/1:20:1:0.2<br>-rm 100:12:0.80:4:0.10:1:0.31:3<br>-rm 100:12:0.80:4:0.15:1:0.31:5<br>-rm 100:13:0.80:4:0.15:1:0.31:4 |

As expected, the BPS results for the HG sampling are higher than those found in Cassava sampling, due to higher entropy of the HG sequence.

All results from HG sampling after 200 generations showed improved compression efficiency but decreased time efficiency compared to the results from the first 100 generations. This may have occurred due to the increased difficulty of the MGA in finding new effective solutions without compromising time and energy resources. The decreased time efficiency was, however, not verified when extending the Cassava sampling method for 200 generations, possibly due to the different strategy used in Cassava sampling compared to HG sampling.

In terms of compression efficiency, the results achieved using HG sampling outperform most of the current entries in the HG benchmark, available at https://github.com/cobilab/HumanGenome, which were previously optimized through exhaustive search.

### 3.7 Human genome sampling experiment (10 MB)

As an extension of the previous experiment, a 10 MB human sample was selected (SG10MB). For this sample, running the MGA for 500 generations, we achieved an impressive compression ratio of 1.431 BPS in the HG benchmark, marking one of the highest compression records to date.

### 3.8 Benchmark results as discussion

The results obtained from the aforementioned optimization experiments were benchmarked by comparing them with those for manually tested JARVIS3 commands and several other state-of-the-art compressors. The benchmark results for CY, Cassava, and HG are illustrated in Fig. 5.

**Figure 5.**
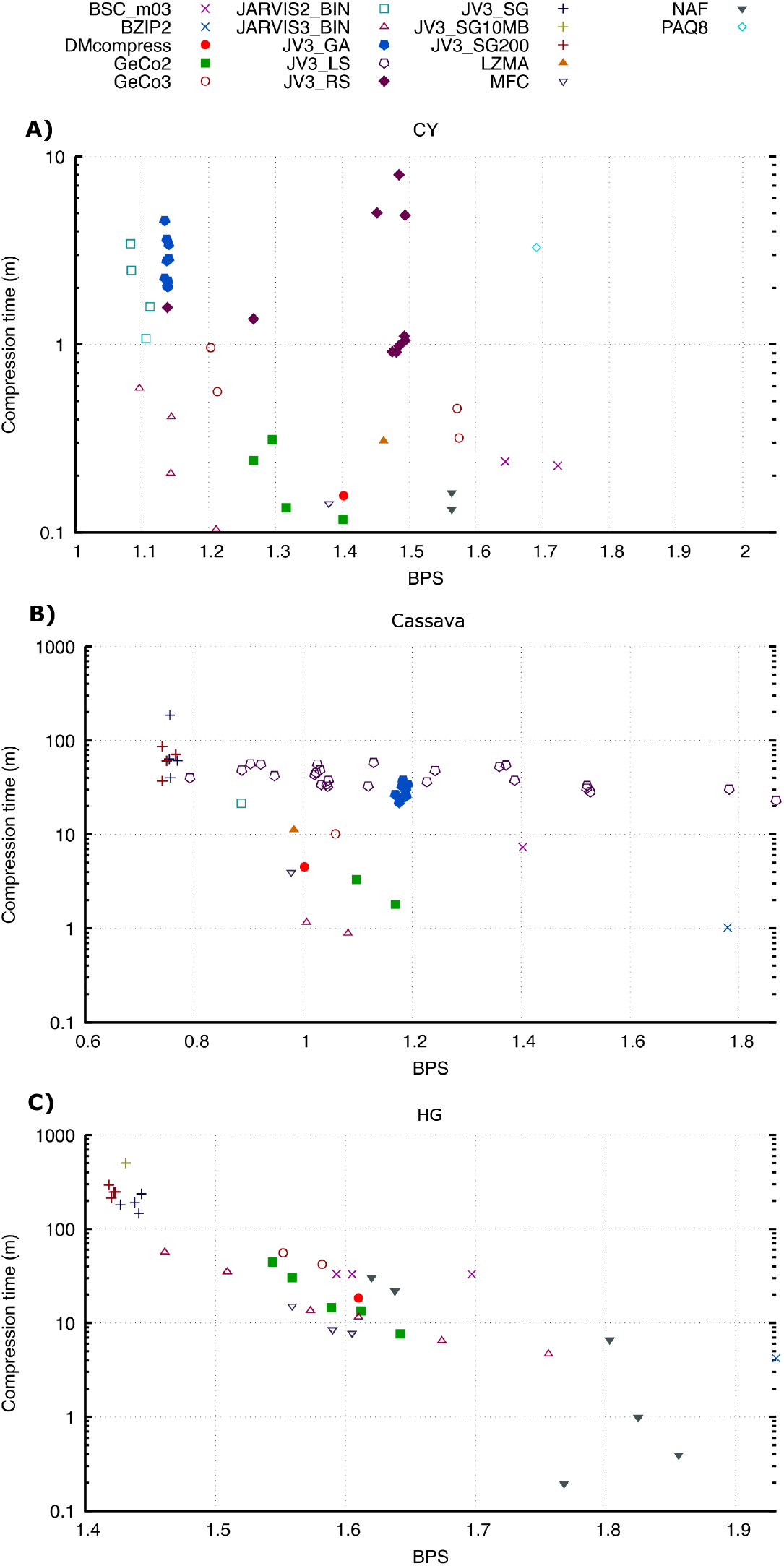
Benchmark results of A) Human chromosome Y (CY); B) Cassava haplotig 1 (Cassava); C) Human genome (HG). The x-axis of each plot corresponds to Bits Per Symbol (BPS).

Although the results found from manually executed tests of JARVIS2 and JARVIS3 are, overall, faster or present better BPS results than those found by OptimJV3, it should be noted that the manually executed tests were more challenging to discover, even with an advanced knowledge in genomic compression. Moreover, it is impractical and inefficient to manually search for the best parameters for a wide range of genomic sequences, due to their varied patterns and lengths. Therefore, the results found from OptimJV3 were overall successful.

The CY plot shows successful BPS results found via MGAs, particularly in compression efficiency. Its BPS results were reasonably similar to those presented by pre-configured tests of JARVIS2 and JARVIS3, though overall, they required a longer time to achieve. The results obtained through RS demonstrate lower performance compared to MGA, highlighting the limitations of RS.

The Cassava plot displays several results found from LS that provide better BPS solutions than those found through MGA. This is unsurprising, considering that the MGA was only executed for 20 generations due to the challenge in applying a MGA to a larger sequence without sampling techniques. Overall, all sampling results found provided better BPS solutions than other methods. However, they were slower than the results obtained through pre-configured JARVIS3 commands.

Similar to the solutions presented in Cassava optimization, the most optimal BPS results for HG compression were achieved through sampling techniques. However, the time efficiency yielded by tests from these techniques was the lowest compared to the other methods explored. To address this issue, the sampling technique could be repeated using a MOGA with a higher weight assigned to time efficiency.

OptimJV3 can now be used in any genomic sequence, making it particularly valuable for optimizing the compression of less-explored yet widely utilized genomes in long-term bioinformatics storage, namely eukaryotes, bacteria, viruses, fungi, and protozoa. This broad applicability not only enhances storage efficiency but also offers insights into the intrinsic complexity of these genomes. Such optimization-driven metrics can serve as informative features in classification tasks, building upon previous studies we conducted without the benefit of optimization, and paving the way for more informed and efficient genomic analysis [38]. Moreover, this framework can now be integrated with genome compressors that support relative compression, such as GeCo3, enabling further gains by leveraging suitable reference sequences during the compression optimization process.

From a practical perspective, OptimJV3 is a one-time search that amortizes: we tune once on a 10–50 MB representative sample for 100–200 generations (or use SOGA with fewer generations on small inputs) and then reuse the discovered JARVIS3 command for the full dataset and across related inputs; the results reported here reflect only the final compression pass, with GA overhead incurred once during tuning. The optimized commands we provide can be adopted directly by end users, avoiding any search in routine use.

Beyond storage efficiency, optimized parameters for widely used reference genomes improve compression-based analyses—yielding stronger detection of rearrangements and structural variants [44], more accurate metagenomic classification [39], improved viral genome reconstruction [45], enhanced temporal analyses of biological sequences [57], and more reliable mapping/flagging of low-complexity regions [58].

For practical use of OptimJV3, we recommend tuning on a representative sample (10-50 MB for large inputs) for 100–200 generations with MOGA (balanced weights), then applying the discovered command to the full genome; small inputs can use SOGA on the full data with fewer generations.

## 4 Conclusions

We developed OptimJV3, an open-source tool that leverages GAs to optimize the JARVIS3 genomic compression model. By efficiently exploring its large parameter space, OptimJV3 consistently improved compression performance across various genomes, including notable gains on Cassava and the HG. The integration of multi-objective optimization enabled simultaneous improvement in compression efficiency and runtime, addressing the pressing need for scalable, lossless compression in genomics.

Among the strategies evaluated, sampling-based optimization was most effective. We tuned parameters on representative subsets, namely 100 MB for Cassava and 25 MB for HG, over 200 generations, then applied the best settings to the full datasets. This yielded 0.741 BPS for Cassava in approximately 37 minutes and 1.418 BPS for HG in approximately 5 hours. These results show that small samples can reliably guide optimization for large-scale sequences.

GA methods also performed well on sequences under 100 MB without sampling. For example, the Human CY achieved 1.134 BPS using modified RWS and 1.139 BPS with MRC, both over 100 generations. MOGAs further showed potential for fast, efficient compression, achieving 1.18 BPS for CY in just 20 seconds. Overall, GAs outperformed non-automated and local search methods, offering a promising solution for efficient, scalable, and potentially more sustainable genomic data compression.

Although JARVIS3 supports both assembled sequences (FASTA) and raw reads (FASTQ) with parallel execution, we deliberately restrict this study to assembled sequences to validate the metameric GA optimizer under controlled conditions and to derive reusable presets for widely used references (CY, Cassava, CHM13/HG). Because such references are repeatedly compressed and widely mirrored, the one-time tuning cost amortizes across extensive reuse. We acknowledge, however, that the absence of read-level (FASTQ) experiments is a limitation of this work, where performance/throughput trade-offs may differ for single-cell, spatial, or metagenomic workloads.

## Data Availability

The source code and data for replication is freely available at https://github.com/cobilab/OptimJV3.

- Project name: OptimJV3
- Project home page: https://github.com/cobilab/OptimJV3
- Operating system(s): Linux
- Programming language: Bash
- Recommended environment: Conda
- License: GNU License
- RRID: SCR_026705
- Biotools: OptimJV3

### List of abbreviations

BPS: bits per symbol
CY: Human chromosome Y
CNGB: China National Genebank
CMGA: Canonical Metameric Genetic Algorithm
CM: Finite-context Model
GA: Genetic Algorithm
HG: Human genome
LS: Local Search
MCC: Metameric Canonical Crossover
MGA: Metameric Genetic Algorithm
MRC: Metameric Random Crossover
NCBI: National Center for Biotechnology Information
RS: Random Search
RWS: Roulette Wheel Selection
SG: Sampling executed for 100 generations
SG10MB: Sampling executed for 10 MB sample
G200: Sampling executed for 200 generations

## Author contributions statement

A.P. and D.P. designed the experiment. R.F. developed the tool and executed data analysis. R.F., A.P. and D.P. analysed and discussed the results. R.F. and D.P. wrote the manuscript and all authors revised the manuscript.

