## Supplementary Material for "Optimizing Genomic Data Compression with Genetic Algorithms"

### **Supplementary material of** Optimizing Genomic Data Compression with Genetic Algorithms”

R. Ferrolho, A. J. Pinho, and D. Pratas

#### 1 Datasets and materials

Table 1 presents all datasets (sequences) used in the experiments, excluding samples.

#### 2 Random and Local Search Results - Supplementary Material

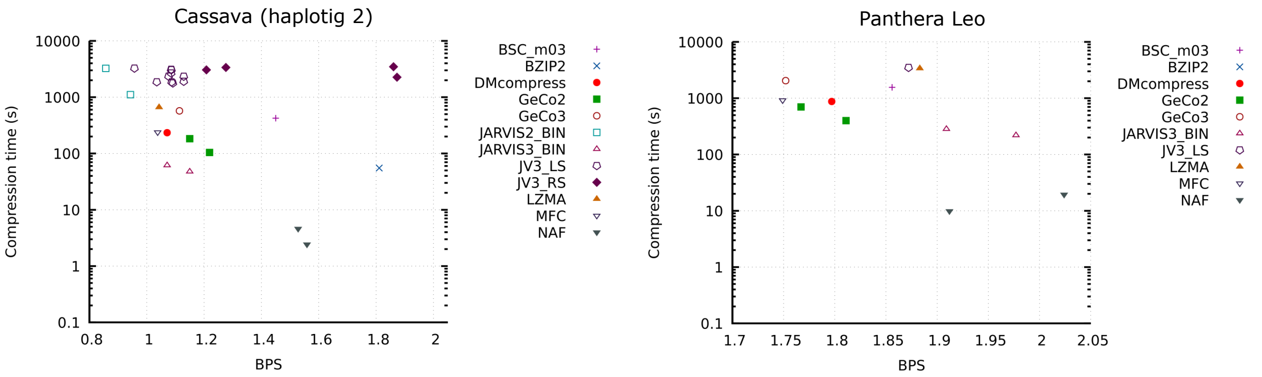


Figure S1: Results of Random Search (RS) for Cassava (haplotig2), and Local Search (LS) for Pathera Leo.

#### 3 Reproducibility

##### 3.1 Quick Demo

To clone the OptimJV3 project and change script permissions, execute the following instructions:

```
git clone https://github.com/cobilab/OptimJV3.git
chmod +x OptimJV3/scripts/*.sh
```

After cloning and activating the script permissions, the following demonstration, where a GA is applied to the compression the human chromosome Y (CY), can be executed in the scripts folder:

```
./GetDSinfo.sh # map sequences into its DS, sorted by size; view sequences info
./GA.sh -s cy -ga "demo" -lg 100 -t 10 # optimize compression of CY using canonical GA for 100
    generations and 10 threads
```

The results will then be stored in a folder with prefix ”DS” within the ”OptimJV3” project. To identify this folder name, the following command can be executed:

```
./Main.sh -v
```

For instance, assuming that there are no other sequences installed, the output folder should be ”DS1”.

##### 3.2 Setup

After cloning and changing script permissions, the following instruction should be executed to install all tools and sequences required to reproduce the experiments:

Table S1: Enumeration of all datasets considered; RS/LS breadth experiments span this full list, whereas MGA/MOGA deep analyses are reported for CY, HG, and Cassava H1/H2. Scripts and outputs for the remaining datasets are available in the repository. ID refers to the NCBI sequence ID or an abbreviation if from another source.

| DS | Source | ID | Description | Size |
|---|---|---|---|---|
| DS1 | NCBI | KT868810.1 | Cutavirus strain BR-283 NS1 gene, partial cds; and putative VP1, hypothetical protein, VP2, and hypothetical protein genes | 4.09668 KB |
| DS2 | NCBI | CM029732.1 | Pollicipes pollicipes isolate AB1234 mitochondrion, complete sequence | 14.7363 KB |
| DS3 | DNACorpus | BuEb | Bundibugyo ebolavirus genome | 18.4961 KB |
| DS4 | NCBI | OM812693.1 | SARS-CoV-2 isolate HUN/Hun-1/2020, complete genome | 29.0693 KB |
| DS5 | AlcoR | alcor30KB | Synthetic sequence | 29.2969 KB |
| DS6 | DNACorpus | AgPh | Aggregatibacter phage S1249 genome | 42.9395 KB |
| DS7 | DNACorpus | YeMi | Yellowstone lake mimivirus genome | 71.9619 KB |
| DS8 | NCBI | NC_001664.4 | Human betaherpesvirus 6A, variant A, isolate U1102, complete genome | 155.643 KB |
| DS9 | NCBI | NC_000898.1 | Human herpesvirus 6B, complete genome | 158.314 KB |
| DS10 | NCBI | NC_000908.2 | Mycoplasmoides genitalium G37, complete sequence | 566.48 KB |
| DS11 | DNACorpus | AeCa | Aeropyrum camini genome | 1.51734 MB |
| DS12 | DNACorpus | HePy | Helicobacter pylori genome | 1.59056 MB |
| DS13 | AlcoR | alcor2MB | Synthetic sequence with five low-complexity regions | 2 MB |
| DS14 | NCBI | NC_004461.1 | Staphylococcus epidermidis ATCC 12228, complete sequence | 2.3835 MB |
| DS15 | DNACorpus | HaHi | Haloarcula hispanica genome | 3.7098 MB |
| DS16 | DNACorpus | EsCo | Escherichia coli genome | 4.42662 MB |
| DS17 | DNACorpus | PlFa | Plasmodium falciparum genome | 8.5704 MB |
| DS18 | DNACorpus | WaMe | Wallemia mellicola genome | 8.72081 MB |
| DS19 | DNACorpus | ScPo | Schizosaccharomyces pombe genome | 10.1587 MB |
| DS20 | NCBI | NC_000024.1 | Human chromosome Y | 21.6181 MB |
| DS21 | DNACorpus | EnIn | Entamoeba invadens genome | 25.1799 MB |
| DS22 | DNACorpus | DrMe | Drosophila miranda chromosome 2 | 30.6906 MB |
| DS23 | NCBI | BA000046.3 | Pan troglodytes DNA, chromosome 22, complete sequence | 31.1909 MB |
| DS24 | NCBI | NC_000021.9 | Human chromosome 21 | 38.2315 MB |
| DS25 | DNACorpus | OrSa | Oriza sativa Japonica chromosome 1 | 41.2584 MB |
| DS26 | NCBI | NC_073246.2 | Gorilla gorilla gorilla isolate KB3781 chromosome 22 | 55.7969 MB |
| DS27 | DNACorpus | DaRe | Danio rerio chromosome 3 | 59.6667 MB |
| DS28 | NCBI | NC_072005.2 | Pongo abelii isolate AG06213 chromosome 20 | 62.3704 MB |
| DS29 | DNACorpus | AnCa | Anolis carolinensis genome | 135.603 MB |
| DS30 | NCBI | NC_000008.11 | Human chromosome 8 | 138.062 MB |
| DS31 | NCBI | NC_058373.1 | Felis Catus chromosome B3 | 141.611 MB |
| DS32 | DNACorpus | GaGa | Gallus gallus chromosome 2 | 141.651 MB |
| DS33 | DNACorpus | HoSa | Human chromosome 4 | 180.962 MB |
| DS34 | CNGB[1] | Cassava2 | Cassava (haplotig 2) | 673.607 MB |
| DS35 | CNGB[2] | Cassava | Cassava (haplotig 1) | 727.074 MB |
| DS36 | NCBI | GCA.008795835.1 | Panthera Leo genome | 2.22454 GB |
| DS37 | [3] | HG | Human genome | 2.9032 GB |

[1] https://s3.ap-northeast-1.wasabisys.com/gigadb-datasets/live/pub/10.5524/102001_103000/102193/00_Assembly_Fasta/haplotigs/TME204.HiFi_HiC.haplotig2.fa
[2] https://s3.ap-northeast-1.wasabisys.com/gigadb-datasets/live/pub/10.5524/102001_103000/102193/00_Assembly_Fasta/haplotigs/TME204.HiFi_HiC.haplotig1.fa
[3] https://s3-us-west-2.amazonaws.com/human-pangenomics/T2T/CHM13/assemblies/analysis_set/chm13v2.0.fa.gz

```
./Setup.sh
```

Alternatively, in the scripts folder, the following instructions should be executed:

```
./InstallTools.sh       # install listed compressors, GTO, and AlcoR
./DownloadFASTA.sh      # downloads FASTA files
./GetCassava.sh         # gunzip cassava files
./GetAlcoRFASTA.sh      # simulates and stores 2 synthetic FASTA sequences
./FASTA2seq.sh          # cleans FASTA files and stores raw sequence files
./DownloadDNAcorpus.sh # download raw sequences from a balanced sequence corpus
./GetSample.sh -s cassava -sp 0.4 -mb 100 -so cassava100MB
./GetSample.sh -s cassava -sp 0.4 -mb 50 -so cassava50MB
./GetSample.sh -s cassava -sp 0.4 -mb 25 -so cassava25MB
./GetSample.sh -s cassava -sp 0.4 -mb 12.5 -so cassava12d5MB
./GetSample.sh -s human -sp 0.2 -mb 10 -so human10MB
./GetDSinfo.sh          # map sequences into their ids, sorted by size; view sequences info
```

##### 3.3 Reproducing the "Non-MGA results"

To reproduce this experiment, the following commands should be executed in the scripts folder:

```
bash -x ./GA.sh -s $sequence -ga "rs" -lr 0 -lg 1 1> out 2> err &
```

##### 3.4 Reproducing the "Canonical MGA experiment"

To reproduce this experiment, the following commands should be executed in the scripts folder:

```
bash -x ./GA.sh -s "escherichia_coli" -ga "e0_ga1_lr0_cmga" -lr 0 -lg 100 1> out 2> err &
bash -x ./GA.sh -s cy -ga "e0_ga1_lr0_cmga" -lr 0 -lg 100 1> out 2> err &
bash -x ./GA.sh -s cassava -ga "e0_ga1_lr0_cmga" -lr 0 -lg 20 1> out 2> err &
```

##### 3.5 Reproducing the "CY experiments"

To reproduce these experiments, the following command should be executed in the scripts folder:

```
bash -x ./Main.sh -s cy -lg 100 -t 10 1> out 2> err &
```

##### 3.6 Reproducing the "Cassava sampling experiment"

To reproduce this experiment, run:

```
./SamplingDemoCassava.sh
```

##### 3.7 Reproducing the "Human genome sampling experiments"

To reproduce this experiment for human genome samples greater than 10 MB, run:

```
./SamplingDemo.sh
```

As for the 10 MB experiment, the following command should be executed to run a GA algorithm for 500 generations:

```
bash -x ./GA.sh -s human10MB -lg 500 1> out 2> err &
```

#### 4 Parameters and options of OptimJV3

All implemented features are listed in the following scripts:

```
./InstallTools.sh -h
./GetDSinfo.sh -h
./Main.sh -h
./GA.sh -h
./Initialization.sh -h
./Run.sh -h
./Evaluation.sh -h
./Selection.sh -h
./CrossMut.sh -h
./GetSamples.sh -h
./PlotGA.sh -h
./PlotGAcmp.sh -h
```

#### 4.1 Install tools menu

To access the menu of the script that installs all required tools, the following command can be executed

```
./InstallTools.sh -h
```

This command will output the following content

```
 ----------------------------------------------------

 OptimJV3 - optimize JARVIS3 CM and RM parameters

 Program options ---------------------------------------

-h|--help.....................................Show this
-iwc|--install-with-conda........Install some tools with
                                                    conda
-iwb|--install-with-both..........Install all tools with
                                        conda and globally

 ----------------------------------------------------
```

with the options/parameters available.

#### 4.2 Download FASTA sequences menu

To access the menu of the script that downloads a set of FASTA sequences, or a specifc FASTA sequence given its NCBI ID, the following command can be executed

```
./DownloadFASTA.sh -h
```

This command will output the following content

```
----------------------------------------------------

CompressSequences - JARIVS3 Optimization Benchmark
Download FASTA files script

Program options ---------------------------------------

--help|-h.....................................Show this
-id........................Download sequence by NCBI id

----------------------------------------------------
```

with the options/parameters available.

#### 4.3 Map sequences to dataset ID menu

The GetDSinfo.sh maps and displays information about sequences, their dataset id, and their size. The command to access its features is:

```
./GetDSinfo.sh -h
```

This command will output the following content

```
 ----------------------------------------------------

 OptimJV3 - optimize JARVIS3 CM and RM parameters

 Program options ---------------------------------------

-h|--help.....................................Show this
-v|--view-ds|--view-datasets.........View datasets info

 ----------------------------------------------------
```

with the options/parameters available.

#### 4.4 Main menu

The main script runs a set of pre-configured GAs and applied to a set of sequences.

The command to access the main menu with its features is

```
./Main.sh -h
```

This command will output the following content

```
---------------------------------------------------------

 OptimJV3 - optimize JARVIS3 CM and RM parameters

 Program options ---------------------------------------

-h|--help......................................Show this
-v|--view-ds|--view-datasets...View sequence names, size
          of each in bytes, MB, and BG, and their group
-s|--seq|--sequence..........Select sequence by its name
-sg|-sequence-grp|--seq-group..Select group of sequences
                                            by their size
-ds|--dataset......Select sequence by its dataset number
-dr|--drange|--dsrange|--dataset-range...........Select
                  sequences by range of dataset numbers
-fg|--first-gen|--first-generation.........Define first
                                       generation number
-lg|--last-gen|--last-generation............Define last
                                       generation number
-rg|--range-gen|--range-generation.....Define generation
                                                   range
-t|--nthreads....Define number of threads to run JARVIS3
                                             in parallel
-sd|--seed.....................Define pseudo-random seed
-si|--seed-increment...............Define seed increment

example 1: ./Main.sh -s human
example 2: ./Main.sh -s cassava -s human

---------------------------------------------------------
```

with the options/parameters available.

#### 4.5 GA menu

The GA script applies a GA to a set of sequences.

The command to access the GA menu with the options/parameters of OptimJV3 is

```
./GA.sh -h
```

This command will output the following content

```
---------------------------------------------------------

 OptimJV3 - optimize JARVIS3 CM and RM parameters

 Program options ---------------------------------------

-h|--help......................................Show this
-v|--view-ds|--view-datasets...View sequence names, size
          of each in bytes, MB, and BG, and their group
-fg|--first-generation...Specify first generation number
-lg|--last-generation......Select last generation number
                                            by their size
-a|-ga|--genetic-algorithm...Define (folder) name of the
                                       genetic algorithm
```

```
-s|--seq|--sequence................Select sequence name
-sg|--seq-grp|--sequence-group.....Select sequence group
-ds|--dataset......Select sequence by its dataset number
-dr|--drange|--dsrange|--dataset-range...........Select
                   sequences by range of dataset numbers
-ps|--psize|--population|--population-size........Define
                                         population size
-sd|--seed....................Define pseudo-random seed
-si|--seed-increment..............Define seed increment

 Program options (initialization) ----------------------

-hei|--heuristic-initialization.......Activate heuristic
                     initialization/local initialization
-hyi|--hybrid-initialization............Activate hybrid
                                          initialization
-hhp|--hybrid-heuristic-percentage.....Define percentage
            of tests generated by heuristic/local search
-hhp|--hybrid-heuristic-percentage........Define number
            of tests generated by heuristic/local search
-mCM|--m-cm|--min-cm....Define minimum number of context
                                            models (CMs)
-MCM|--M-cm|--max-cm...........Define maximum number of
                                    context models (CMs)
-mRM|--m-rm|--min-rm...........Define minimum number of
                                copy/repeat models (RMs)
-MRM|--M-rm|--max-rm...........Define maximum number of
                                copy/repeat models (RMs)
-lr|--learning-rate................Define learning rate
-hs|--hidden-size....................Define hidden size
-sing|--seeding................Activate seeding feature
to populate with few hardcoded solutions. Only works for
                                           human genome

 Program options (run) ---------------------------------

-t|--nthreads....Define number of threads to run JARVIS3
                                             in parallel

 Program options (evaluation) --------------------------

--moga|--moga-wm|--moga-weighted-metric...Activate Multi
       Objective Genetic Algorithm (MOGA) using weighted
                                         metric function
--moga-ws|--moga-weighted-sum...Activate Multi-Objective
    Genetic Algorithm (MOGA) using weighted sum function
-pe|--p-exp|--p-expoent..........Define expoent for MOGA
                                weighted metric function
-w1|-wBPS|--w-bps|--weight-bps.....Bits Per Symbol (BPS)
                                weight for MOGA function
-w2|-wCTIME|--w-ctime|--weight-ctime....Compression time
                                weight for MOGA function

 Program options (selection) ---------------------------

-sl|--sel|--selection..................Choose selection
   operator: 'elitist' (default), 'roulette', 'rank', or
                                            'tournament'
-ns|--num-sel-cmds...Define number of commands to select
-sr|--selection-rate...Define rate of commands to select

```

```
 Program options (crossover) --------------------------

-cr|-ccr|--command-crossover-rate..Define crossover rate
                            of a selected pair of commands
-mrc|--model-crossover-rate.......Define crossover rate
                          of a selected pair of CMs or RMs
-cc|--command-crossover.........Choose command crossover
 operator: 'mrc' (metameric random crossover) (default),
                     (metameric canonical crossover) 'mcc'
-c|--crossover..........Choose model crossover operator:
   'xpoint' (default), 'uniform', 'average', 'discrete',
                                   'flat', 'heuristic'

 Program options (mutation) ---------------------------

-hm|--heuristic-mutation.....Activate heuristic mutation
                              for narrower range mutations
-mr|-cmr|--command-mutation-rate....Define mutation rate
                                            of a command
-pmr|--parameter-mutation-rate......Define mutation rate
                                          of a parameter

Program options (stop criteria) ------------------------

-sc|--stop-criteria.........Define stop criteria: '1' to
             halt program when no offspring is produced,
                  else (default) stop at last generation
```

with the options/parameters available.

#### 4.6 Initialization menu

The initialization script starts a GA by initializing a population of commands that compress a chosen sequence.

To access the initialization menu, the following menu can be executed

```
./Initialization.sh -h
```

This command will output the following content

```
 -------------------------------------------------------

 OptimJV3 - optimize JARVIS3 CM and RM parameters

 Program options ---------------------------------------

 -h|--help....................................Show this
 -v|--view-ds|--view-datasets..View sequence names, size
           of each in bytes, MB, and BG, and their group
-s|--seq|--sequence..........Select sequence by its name
-sg|--sequence-grp|--seq-group.Select group of sequences
                                           by their size
-a|-ga|--genetic-algorithm...Define (folder) name of the
                                       genetic algorithm
-s|--seq|--sequence.................Select sequence name
-sg|--seq-grp|--sequence-group.....Select sequence group
-ds|--dataset......Select sequence by its dataset number
-dr|--drange|--dsrange|--dataset-range...........Select
                     sequences by range of dataset numbers
-ps|--psize|--population|--population-size.......Define
                                         population size
-sd|--seed.....................Define pseudo-random seed
-si|--seed-increment...............Define seed increment

```

```
 Program options (initialization) ---------------------

-hei|--heuristic-initialization.......Activate heuristic
                      initialization/local initialization
-hyi|--hybrid-initialization.............Activate hybrid
                                           initialization
-hhp|--hybrid-heuristic-percentage.....Define percentage
            of tests generated by heuristic/local search
-hhp|--hybrid-heuristic-percentage.........Define number
            of tests generated by heuristic/local search
-mCM|--m-cm|--min-cm....Define minimum number of context
                                             models (CMs)
-MCM|--M-cm|--max-cm...........Define maximum number of
                                     context models (CMs)
-mRM|--m-rm|--min-rm...........Define minimum number of
                                 copy/repeat models (RMs)
-MRM|--M-rm|--max-rm...........Define maximum number of
                                 copy/repeat models (RMs)
-lr|--learning-rate................Define learning rate
-hs|--hidden-size....................Define hidden size
-sing|--seeding................Activate seeding feature
to populate with few hardcoded solutions. Only works for
                                             human genome
```

with the options/parameters available.

#### 4.7 Run menu

The run script is used to run the initial population or offspring from a certain generation.

To access the menu of the run script, the following command can be executed

```
./Run.sh -h
```

This command will output the following content

```
 ---------------------------------------------------------

 OptimJV3 - optimize JARVIS3 CM and RM parameters

 Program options ----------------------------------------

-h|--help....................................Show this
-v|--view-ds|--view-datasets...View sequence names, size
           of each in bytes, MB, and BG, and their group
-s|--seq|--sequence..........Select sequence by its name
-sg|--sequence-grp|--seq-group.Select group of sequences
                                            by their size
-a|-ga|--genetic-algorithm...Define (folder) name of the
                                        genetic algorithm
-ds|--dataset......Select sequence by its dataset number
-dr|--drange|--dsrange|--dataset-range...........Select
                    sequences by range of dataset numbers
-ps|--psize|--population|--population-size........Define
                                          population size
-sd|--seed.....................Define pseudo-random seed
-si|--seed-increment..............Define seed increment

 Program options (run) --------------------------------

-t|--nthreads....Define number of threads to run JARVIS3
                                              in parallel
-to|--timeout..........Define timeout (default: 1 hour)
```

with the options/parameters available.

#### 4.8 Evaluation menu

This script is used to evaluate and sort the current generation based on their compression, expressed in BPS, and compression time.

To access the menu of evaluation script, the following command can be executed

```
./Evaluation.sh -h
```

This command will output the following content

```
 -------------------------------------------------------

 OptimJV3 - optimize JARVIS3 CM and RM parameters

 Program options ---------------------------------------

-h|--help......................................Show this
-v|--view-ds|--view-datasets...View sequence names, size
           of each in bytes, MB, and BG, and their group
-s|--seq|--sequence..........Select sequence by its name
-sg|--sequence-grp|--seq-group.Select group of sequences
                                           by their size
-a|-ga|--genetic-algorithm...Define (folder) name of the
                                       genetic algorithm
-s|--seq|--sequence.................Select sequence name
-sg|--seq-grp|--sequence-group.....Select sequence group
-ds|--dataset......Select sequence by its dataset number
-dr|--drange|--dsrange|--dataset-range............Select
                   sequences by range of dataset numbers
-ps|--psize|--population|--population-size........Define
                                         population size
-sd|--seed.....................Define pseudo-random seed
-si|--seed-increment...............Define seed increment

 Program options (evaluation) --------------------------

--moga|--moga-wm|--moga-weighted-metric...Activate Multi
       Objective Genetic Algorithm (MOGA) using weighted
                                         metric function
--moga-ws|--moga-weighted-sum...Activate Multi-Objective
    Genetic Algorithm (MOGA) using weighted sum function
-pe|--p-exp|--p-expoent..........Define expoent for MOGA
                                weighted metric function
-w1|-wBPS|--w-bps|--weight-bps.....Bits Per Symbol (BPS)
                                weight for MOGA function
-w2|-wCTIME|--w-ctime|--weight-ctime....Compression time
                                weight for MOGA function
```

with the options/parameters available.

#### 4.9 Selection menu

This script selects commands from the population as potential candidates for producing offspring.

To access the selection menu, the following command can be executed

```
./Selection.sh -h
```

This command will output the following content

```
 -------------------------------------------------------

 OptimJV3 - optimize JARVIS3 CM and RM parameters

 Program options ---------------------------------------

```

```
-h|--help....................................Show this
-v|--view-ds|--view-datasets...View sequence names, size
          of each in bytes, MB, and BG, and their group
-s|--seq|--sequence..........Select sequence by its name
-sg|--sequence-grp|--seq-group.Select group of sequences
                                            by their size
-a|-ga|--genetic-algorithm...Define (folder) name of the
                                        genetic algorithm
-s|--seq|--sequence................Select sequence name
-sg|--seq-grp|--sequence-group.....Select sequence group
-ds|--dataset......Select sequence by its dataset number
-dr|--drange|--dsrange|--dataset-range...........Select
                  sequences by range of dataset numbers
-ps|--psize|--population|--population-size.......Define
                                          population size
-sd|--seed....................Define pseudo-random seed
-si|--seed-increment..............Define seed increment

 Program options (selection) ---------------------------

-sl|--sel|--selection..................Choose selection
   operator: 'elitist' (default), 'roulette', 'rank', or
                                             'tournament'
-ns|--num-sel-cmds...Define number of commands to select
-sr|--selection-rate...Define rate of commands to select
```

with the options/parameters available.

##### 4.10 Crossover and Mutation menu

The crossover and mutation script is used to produce offspring.

To access the menu of this script, the following command can be executed

```
./CrossMut.sh -h
```

This command will output the following content

```
 ------------------------------------------------------

 OptimJV3 - optimize JARVIS3 CM and RM parameters

 Program options ---------------------------------------

-h|--help....................................Show this
-v|--view-ds|--view-datasets...View sequence names, size
          of each in bytes, MB, and BG, and their group
-s|--seq|--sequence..........Select sequence by its name
-sg|--sequence-grp|--seq-group.Select group of sequences
                                            by their size
-a|-ga|--genetic-algorithm...Define (folder) name of the
                                        genetic algorithm
-s|--seq|--sequence................Select sequence name
-sg|--seq-grp|--sequence-group.....Select sequence group
-ds|--dataset......Select sequence by its dataset number
-dr|--drange|--dsrange|--dataset-range...........Select
                  sequences by range of dataset numbers
-ps|--psize|--population|--population-size.......Define
                                          population size
-sd|--seed....................Define pseudo-random seed
-si|--seed-increment..............Define seed increment

 Program options (crossover) ---------------------------

```

```
-cr|-ccr|--command-crossover-rate..Define crossover rate
                              of a selected pair of commands
-mrc|--model-crossover-rate........Define crossover rate
                            of a selected pair of CMs or RMs
-cc|--command-crossover.........Choose command crossover
 operator: 'mrc' (metameric random crossover) (default),
                     (metameric canonical crossover) 'mcc'
-c|--crossover..........Choose model crossover operator:
   'xpoint' (default), 'uniform', 'average', 'discrete',
                                     'flat', 'heuristic'

 Program options (mutation) ----------------------------

-hm|--heuristic-mutation.....Activate heuristic mutation
                                for narrower range mutations
-mr|-cmr|--command-mutation-rate....Define mutation rate
                                                of a command
-pmr|--parameter-mutation-rate......Define mutation rate
                                              of a parameter
```

with the options/parameters available.

##### 4.11 Get Sample menu

The features of the script that outputs a sample file can be found by running the following command

```
./GetSample.sh -h
```

This command will output the following content

```
 -------------------------------------------------------

 OptimJV3 - optimize JARVIS3 CM and RM parameters

 Program options ---------------------------------------

-h|--help.....................................Show this
-s|--sequence..................Choose sequence name/file
-sp|--start-percentage.....Define Percentage of the full
                                sequence where sample begins
-sz|-mb|--size-mb....................Define sample size
-so|--sample-output........Define sample output filename

 -------------------------------------------------------

```

with the options/parameters available.

##### 4.12 Get Samples menu

The features of the script that outputs four sample files from a sequence - with sizes of 100MB, 50MB, 25MB and 12.5MB - can be found by running the following command

```
./GetSamples.sh -h
```

This command will output the following content

```
 -------------------------------------------------------

 OptimJV3 - optimize JARVIS3 CM and RM parameters

 Program options ---------------------------------------

-h|--help.....................................Show this
-s|--seq|--sequence............Choose sequence name/file
```

```

-------------------------------------------------------
```

with the options/parameters available.

#### 4.13 Plot GA menu

The features of the script that plots the evolutionary plots and histograms of a single GA can be found by running the following command

```
./PlotGA.sh -h
```

This command will output the following content

```
-------------------------------------------------------

OptimJV3 - optimize JARVIS3 CM and RM parameters

Program options ---------------------------------------

-h|--help......................................Show this
-a|-ga|--genetic-algorithm...Define (folder) name of the
                                       genetic algorithm
-s|--seq|--sequence............Choose sequence name/file
-ds|--dataset......Select sequence by its dataset number
-pb|--percentage-best..........Define percentage of best
                                     individuals to plot
-b|--best......Define number of best individuals to plot
-fg|--first-generation...Specify first generation number
-lg|--last-generation......Select last generation number
-br|--b-range..................Define x-axis (BPS range)
-trs|--trange-s...Define y-axis (time range, in seconds)
-trm|--trange-m...Define y-axis (time range, in minutes)
-trh|--trange-h.....Define y-axis (time range, in hours)
-hi|--hist-interval........Define bin size for histogram

-------------------------------------------------------
```

with the options/parameters available.

#### 4.14 Plot GA comparison menu

The features of the script that plots the comparison between GAs of a specific experiment can be found by running the following command

```
./PlotGAcmp.sh -h
```

This command will output the following content

```
   -------------------------------------------------------

   OptimJV3 - optimize JARVIS3 CM and RM parameters

   Program options ---------------------------------------

  -h|--help......................................Show this
  -e|--experiment-number.........Define experiment number:
       1. Compare GAs for different initialization methods
       2. Compare GAs for different population sizes
       3. Compare MOGAs
       4. Compare GAs for different selection methods
       5. Compare GAs for different crossover methods
  -ds|--dataset......Select sequence by its dataset number
  -fg|--first-generation...Specify first generation number
  -lg|--last-generation......Select last generation number
```

```
  -br|--b-range.................Define x-axis (BPS range)
  -trs|--trange-s...Define y-axis (time range, in seconds)
  -trm|--trange-m...Define y-axis (time range, in minutes)
  -trh|--trange-h.....Define y-axis (time range, in hours)

   -----------------------------------------------------
```

with the options/parameters available.

### 5 Parameters and options of CompressSequences

```
./InstallTools.sh -h
./DownloadFASTA.sh -h
./GetDSinfo.sh -h
./RunTestsExample.sh
./ProcessBenchRes.sh -h
./Plot.sh -h
```

#### 5.1 Install tools menu

See “‘Install tools menu”’ subsection from “‘Parameters and options of OptimJV3”’ section.

#### 5.2 Download FASTA sequences menu

See “‘Download FASTA sequences menu”’ subsection from “‘Parameters and options of OptimJV3”’ section.

#### 5.3 Map sequences to dataset ID menu

See “‘Map sequences to dataset ID menu”’ subsection from “‘Parameters and options of OptimJV3”’ section.

#### 5.4 Plot menu

The features of the script that plots the results of benchmark data can be found by running

```
./Plot.sh -h
```

This command will output the following content

```
   -----------------------------------------------------

   CompressSequences - benchmark
   Plot Script

   Program options ---------------------------------------

   -s|--sequence..........................Select sequence
                                                  of each
  --sequence|--seq|-s.........Select sequence by its name
  --sequence-group|--seq-grp|-sg.Select group of sequences
                                            by their size
              1. Sequences with size lower than 1MB
              2. Sequences with size between 1MB and 100MB
              3. Sequences with size between 100MB and 1GB
              4. Sequences with size between 1GB and 3GB
              5. Sequences with size greater than 3GB
  -br|--b-range.................Define x-axis (BPS range)
  -trs|--trange-s...Define y-axis (time range, in seconds)
  -trm|--trange-m...Define y-axis (time range, in minutes)
  -trh|--trange-h.....Define y-axis (time range, in hours)
  -m|--mode....Select data to plot ('bench', 'NGA', 'all')

   -----------------------------------------------------
```

with the options/parameters available.

#### 5.5 Run menu

The features of the script that runs the generated JARVIS3 commands can be found by running

```
./Run.sh -h
```

Alternatively, to access the menu of a similar script which runs fewer tests the following instruction can be executed:

```
./RunLess.sh -h
```

Both instructions will output the following content

```
 -----------------------------------------------------

 CompressSequences - benchmark
 Run Script

 Program options -------------------------------------

-h|--help....................................Show this
-v|--view-ds|--view-datasets...View sequence names, size
           of each in bytes, MB, and GB, and their group
-s|--seq|--sequence.........Select sequence by its name
-sg|--sequence-grp|--seq-group.Select group of sequences
                                            by their size
-a|-ga|--genetic-algorithm...Define (folder) name of the
                                        genetic algorithm
-ds|--dataset......Select sequence by its dataset number
-dr|--drange|--dsrange|--dataset-range...........Select
                     sequences by range of dataset numbers
-g|--gen-num...................Define generation number
-to|--timeout...........................Define timeout
-t|--nthreads....Define number of threads to run JARVIS3
                                              in parallel
```

with the options/parameters available.

#### 5.6 Process Benchmark results menu

The features of the script that processes the benchmark data can be found by running

```
./ProcessBenchRes.sh -h
```

This command will output the following content

```
  -----------------------------------------------------

  OptimJV3 - optimize JARVIS3 CM and RM parameters

  Program options -------------------------------------

  -h|--help....................................Show this
  -v|--view-ds|--view-datasets....View sequences and size
                                                  of each

  -----------------------------------------------------
```

with the options/parameters available.

#### 5.7 JARVIS3 Parameters

JARVIS3 [1] Context Models (-cm) and Repeat Models (-rm) are the following:

```

      -cm [NB_C]:[NB_D]:[NB_I]:[NB_G]/[NB_S]:[NB_E]:[NB_R]:[NB_A]
      Template of a context model.
      Parameters:
      [NB_C]: (integer [1;14]) order size of the regular context
              model. Higher values use more RAM but, usually, are
              related to a better compression score.
      [NB_D]: (integer [1;5000]) denominator to build alpha, which
              is a parameter estimator. Alpha is given by 1/[NB_D].
              Higher values are usually used with higher [NB_C],
              and related to confident bets. When [NB_D] is one,
              the probabilities assume a Laplacian distribution.
      [NB_I]: (integer {0,1,2}) number to define if a sub-program
              which addresses the specific properties of DNA
```

```
                sequences (Inverted repeats) is used or not. The
                number 1 turns ON the sub-program using at the same
                time the regular context model. The number 2 does
                only contemple the inversions only (NO regular). The
                number 0 does not contemple its use (Inverted repeats
                OFF). The use of this sub-program increases the
                necessary time to compress but it does not affect the
                RAM.
        [NB_G]: (real [0;1)) real number to define gamma. This value
                represents the decayment forgetting factor of the
                regular context model in definition.
        [NB_S]: (integer [0;20]) maximum number of editions allowed
                to use a substitutional tolerant model with the same
                memory model of the regular context model with
                order size equal to [NB_C]. The value 0 stands for
                turning the tolerant context model off. When the
                model is on, it pauses when the number of editions
                is higher that [NB_C], while it is turned on when
                a complete match of size [NB_C] is seen again. This
                is probabilistic-algorithmic model very useful to
                handle the high substitutional nature of genomic
                sequences. When [NB_S] > 0, the compressor used more
                processing time, but uses the same RAM and, usually,
                achieves a substantial higher compression ratio. The
                impact of this model is usually only noticed for
                higher [NB_C].
        [NB_R]: (integer {0,1}) number to define if a sub-program
                which addresses the specific properties of DNA
                sequences (Inverted repeats) is used or not. It is
                similar to the [NR_I] but for tolerant models.
        [NB_E]: (integer [1;5000]) denominator to build alpha for
                substitutional tolerant context model. It is
                analogous to [NB_D], however to be only used in the
                probabilistic model for computing the statistics of
                the substitutional tolerant context model.
        [NB_A]: (real [0;1)) real number to define gamma. This value
                represents the decayment forgetting factor of the
                substitutional tolerant context model in definition.
                Its definition and use is analogus to [NB_G].

        ... (you may use several context models)


        -rm [NB_R]:[NB_C]:[NB_B]:[NB_L]:[NB_G]:[NB_I]:[NB_W]:[NB_Y]
        Template of a repeat model.
        Parameters:
        [NB_R]: (integer [1;10000] maximum number of repeat models
                for the class. On very repetive sequences the RAM
                increases along with this value, however it also
                improves the compression capability.
        [NB_C]: (integer [1;14]) order size of the repeat context
                model. Higher values use more RAM but, usually, are
                related to a better compression score.
        [NB_B]: (real (0;1]) beta is a real value, which is a
                parameter for discarding or maintaining a certain
                repeat model.
        [NB_L]: (integer (1;20]) a limit threshold to play with
                [NB_B]. It accepts or not a certain repeat model.
        [NB_G]: (real [0;1)) real number to define gamma. This value
                represents the decayment forgetting factor of the
                regular context model in definition.
        [NB_I]: (integer {0,1,2}) number to define if a sub-program
                which addresses the specific properties of DNA
                sequences (Inverted repeats) is used or not. The
                number 1 turns ON the sub-program using at the same
                time the regular context model. The number 0 does
                not contemple its use (Inverted repeats OFF). The
                number 2 uses exclusively Inverted repeats. The
                use of this sub-program increases the necessary time
                to compress but it does not affect the RAM.
        [NB_W]: (real (0;1)) initial weight for the repeat class.
        [NB_Y]: (integer {0}, [1;50]) maximum cache size. This will
                use a table cache with the specified size. The size
                must be in balance with the k-mer size [NB_C].
```
